# The Janus face of Selinexor (KPT-330) in bacterial infection: short-term protection versus long-term lethality

**DOI:** 10.1101/2025.10.16.682758

**Authors:** Qiaohua Yan, Lu Guo, Qingxiang Sun

## Abstract

The clinical application of the anti-cancer drug KPT-330 (Selinexor) has been associated with increased infection-related mortality, yet its direct impact on infection pathogenesis remains unclear. Here, we found that KPT-330 treatment reduced host survival from 66% to 22% in a mouse model of bacterial pneumonia. Interestingly, KPT-330 conferred transient lung protection at 24 h post infection by blocking the nuclear export of IκB, suppressing NFκB signaling and immune cell recruitment. Meanwhile, KPT-330 induced apoptosis of lung epithelial cells, enhancing bacterial adhesions and invasion, and leading to dramatically increased bacterial load. These effects culminated in a compensatory immune rebound and more severe lung injury at later stages. Adjunctive therapy with the antibiotic polymyxin E (colistin) rescued survival, whereas the immunosuppressant dexamethasone did not, underscoring that timely bacterial clearance is critical for managing this adverse effect. Our findings provide direct evidence that KPT-330 exacerbates bacterial infection and advocate for an adjunctive antimicrobial prophylaxis strategy to ensure its safer use, especially in high-risk patients.

## Introduction

KPT-330 (selinexor) is a first-in-class, orally available anticancer drug that acts by potently inhibiting the nuclear export factor Exportin 1 (XPO1, CRM1).^1-3^ Its pharmacologic effect lasts 2-3 days, and is administered once or twice per week.^4-6^ Besides being a valuable anticancer therapeutic, KPT-330 has been investigated preclinically for the treatment of several other diseases such as sepsis, alzheimer’s disease, autoimmune diseases.^7-9^ These effects are primarily due to KPT-330’s potent inhibition of the NFκB signaling pathway, a master regulator of inflammation.^9-11^

However, inflammation serves as the body’s essential first-line defense against bacterial pathogens.^12^ Correspondingly, clinical trials of KPT-330 reported frequent serious infectious complications including pneumonia and sepsis, with grade 3-4 events reaching over 20% in some studies.^13-15^ Despite this clear clinical association, the dynamic and direct impact of KPT-330 on the host-pathogen interaction throughout the course of a bacterial infection and the underlying mechanisms remain entirely unexplored.

In this work, we employed a murine model of *Acinetobacter baumannii* (Ab) pneumonia to dissect this complex relationship. Our results revealed a dualistic, time-dependent role for KPT-330: it provided an initial, short-term protection against acute lung damage by suppressing NFκB signaling and inflammation, but ultimately induced epithelial cell apoptosis and enhanced bacterial adhesion and invasion, leading to an uprise in bacterial load, tissue pathology, and host lethality at later stages. Lastly, we showed that the side effect of KPT-330 can be successfully reversed by adjunct antibiotic therapy but not by immunosuppressant therapy.

## Results

### KPT-330 increases the bacterial load of infected mice and reduces survival

Twenty-four hours prior to Ab injection, mice were precondition with 20 mg/kg KPT-330 or isotype control (Fig. 1A). Mice were harvested after at 24 h, 48 h, and 72 h post infection (n=5 each). In KPT-330-treated mice, the bacterial load in the lung increased approximately 80-fold at 24 h and 23-fold at 48 h (Fig. 1B). By 72 h, Ab was largely eliminated in both groups. Bronchoalveolar lavage fluid (BALF) also displayed a higher bacterial load in KPT-330-treated mice at 24 h and 48 h (Fig. 1C). In addition, bacterial loads in the blood, liver, spleen, and kidney were elevated in the KPT-330-treated group at 24 h and 48 h, although these differences did not reach statistical significance (Fig. 1D-1G).

**Fig. 1.**
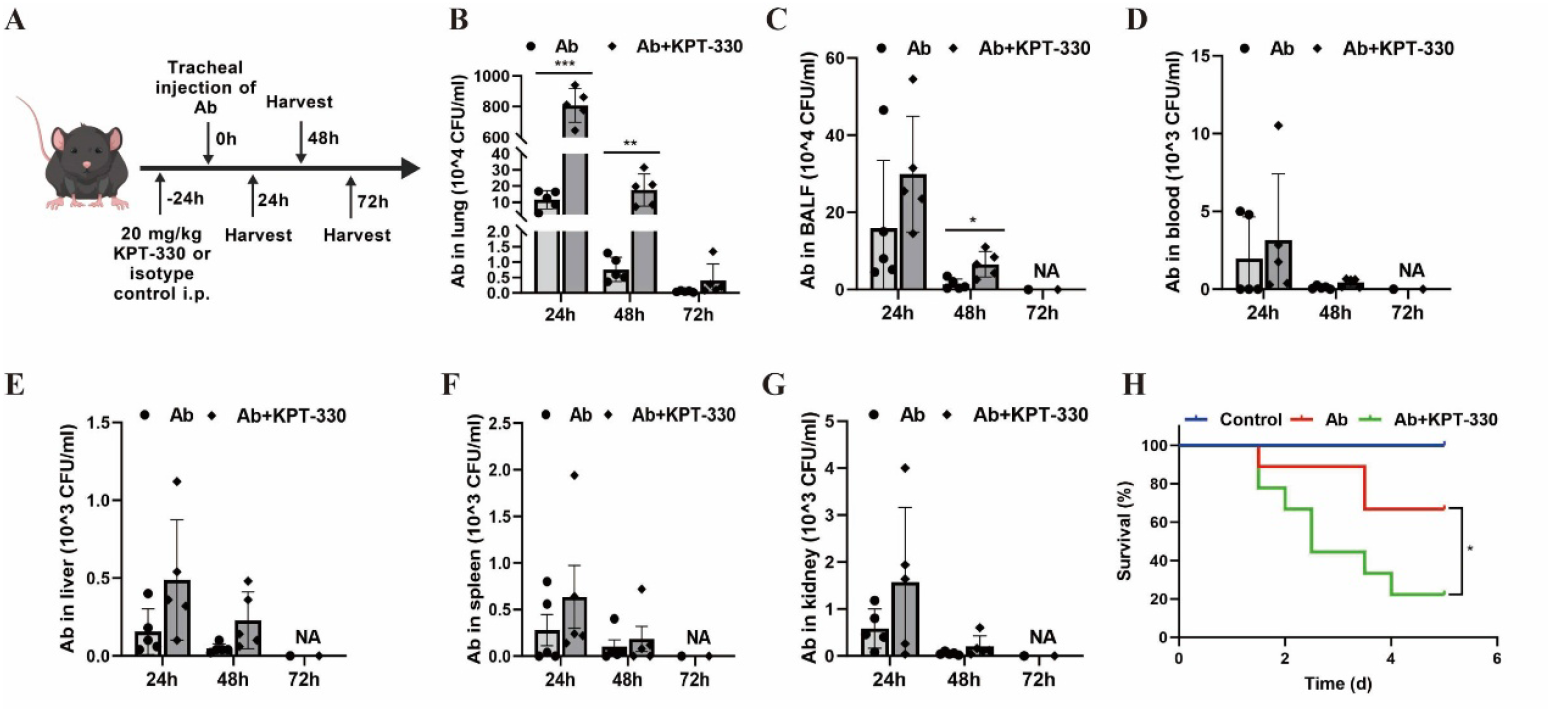
KPT-330 increases the bacterial load of infected mice and reduces survival. (A) Design diagram of the mice experiment plan. Mice were intraperitoneally injected with KPT-330 (20 mg/kg) 24 h prior to intratracheal instillation of Ab (8×10^7^ CFU) to induce acute pneumonia. Lung tissues and BALF were collected at 24 h, 48 h, and 72 h post-induction (n=5 each). (B-G) Bacterial loads in lung tissues, BALF, blood, liver, spleen, and kidney at 24 h, 48 h, and 72 h post-induction. CFU, colony forming units. (H) The survival rates of mice (n=9 each) instilled with 1×10^8^ CFU Ab and treated with or without KPT-330 (20 mg/kg). Data are presented as mean ± standard deviations (SD). Statistical significance was determined using unpaired Student’s t test. *p < 0.05, **p < 0.01, ***p < 0.001.

We further analyzed the survival rates of KPT-330- or iso-type-treated mice infected with a slightly higher dose of Ab for 5 days, with 9 mice in each group. KPT-330 treatment dramatically reduced the survival rate from 66% to 22% at day 5 (Fig. 1H, p < 0.05). These results agree with the immune-suppressive role of KPT-330 and confirm that KPT-330 exacerbates bacterial infection and reduces survival under certain circumstances like Ab pneumonia.

### KPT-330-treated mice exerts a biphasic effect on lung injury

To explore the mechanism underlying KPT-330-induced mortality, we performed histological analysis on lungs from each group. Representative histological images reveals that Ab alone induced lung damage at 24 h, as evidenced by collapsed alveolar spaces, the exudation of red blood cells or fibrin into the alveolar lumens, and swollen capillary endothelial cells (Fig. 2A-2C). These defects were gradually and partially resolved at 48 h and 72 h (Fig. 2A-2C). Interestingly, KPT-330 treatment ameliorated lung damage at 24 h, but caused significantly more severe damage by 72 h, compared to the Ab-only group. In addition, we observed that KPT-330 significantly reduced lung index (lung weight normalized to body weight) and wet-to-dry (W/D) ratio of lung at 24 h (p < 0.01 and p < 0.05), while it slightly increased these values at 72 h (Fig. 2D and 2E). Furthermore, total protein levels in the BALF, which partly reflect lung damage, were significantly suppressed at 24 h and elevated at 48 h and 72 h in KPT-330-treated mice (Fig. 2F). The level of lactate dehydrogenase, a marker of cell membrane leakage, was likewise initially reduced and subsequently enhanced (Fig. S1). These results suggest that KPT-330 provides transient protection against acute lung damage but ultimately exacerbates tissue injury.

**Fig. 2.**
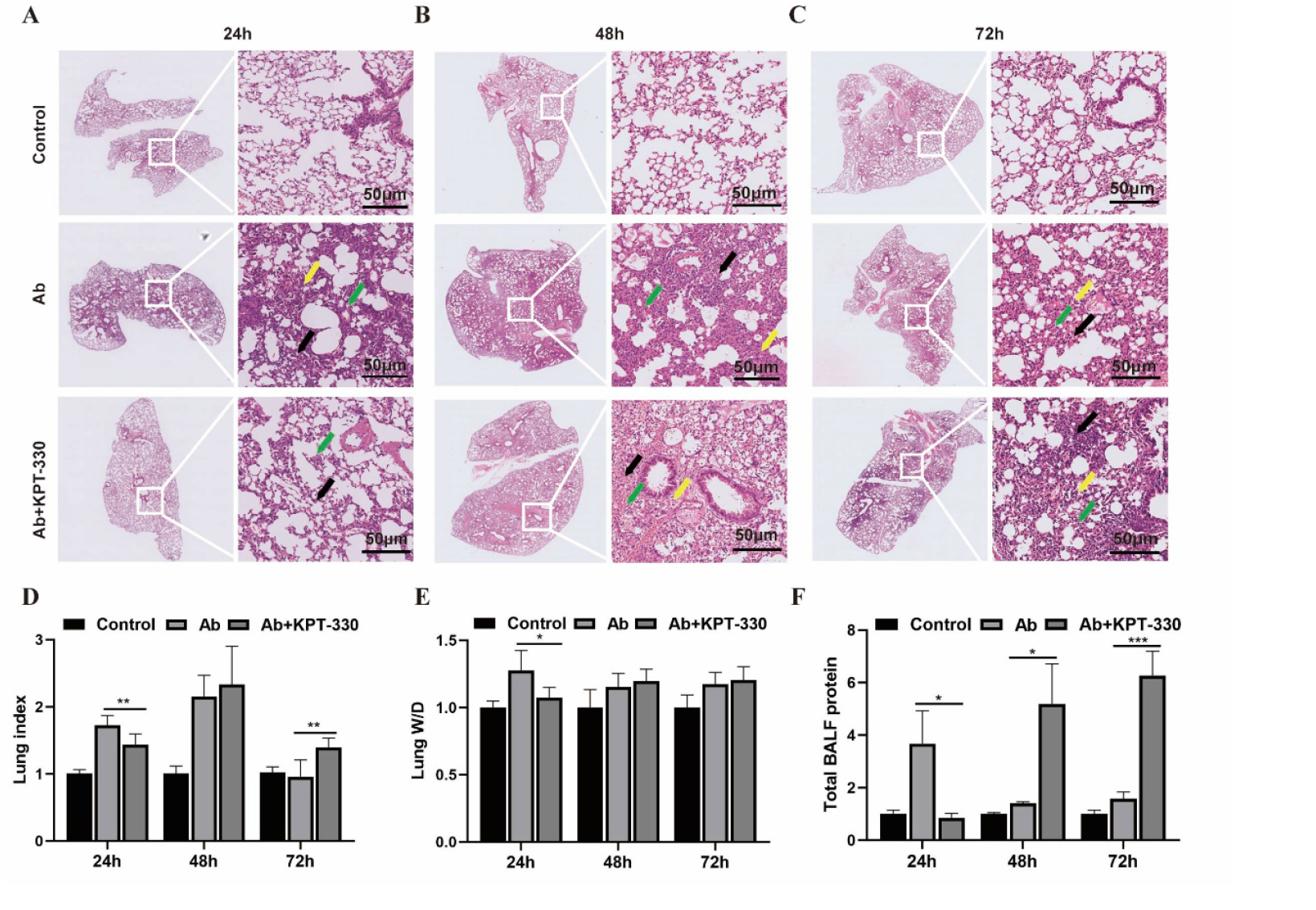
KPT-330-treated mice exerts a biphasic effect on lung injury. (A-C) Representative H&E staining of mice lung tissues at 24 h, 48 h, and 72 h post-induction. Black arrows, thickened alveolar walls accompanied by collapsed alveolar spaces; yellow arrows, red blood cells or fibrin exudation in alveolar lumens; green arrows, swollen capillary endothelial cells. Scale bar, 50 μM. (D) The lung index of mice calculated as lung/body weight. (E) The wet-to-dry (W/D) ratio of the lungs of mice at 24 h, 48 h, and 72 h. (F) The total protein content in the BALF of mice determined using the bicinchoninic acid (BCA) method. All data were normalized to control (n=5 each). Data are presented as mean ± SD. Differences between groups were compared using unpaired Student’s t test. *p < 0.05, **p < 0.01, ***p < 0.001.

#### KPT-330 induces epithelial cell apoptosis by promoting bacterial adhesion and invasion

In agreement with the known effect of KPT-330 inducing apoptosis in cancer cells,^16-18^ it was observed that KPT-330-treated lung displayed the highest apoptosis level regardless of time, as measured by the ratio of proapoptotic factor Bax and anti-apoptotic factor Bcl-2 (Fig. 3A). In Ab-treated lung epithelial Beas-2b cells, KPT-330 also increased the of Bax/Bcl-2 ratio, PARP-1 cleavage, while decreasing Bcl-XL levels (Fig. 3B). In contrast, Ab alone showed minimal effect on these pro/anti-apoptotic factors or the cleavage of PARP-1. Flow cytometry analysis further corroborated the pro-apoptotic role of KPT-330 in Beas-2b cells (Fig. 3C).

**Fig. 3.**
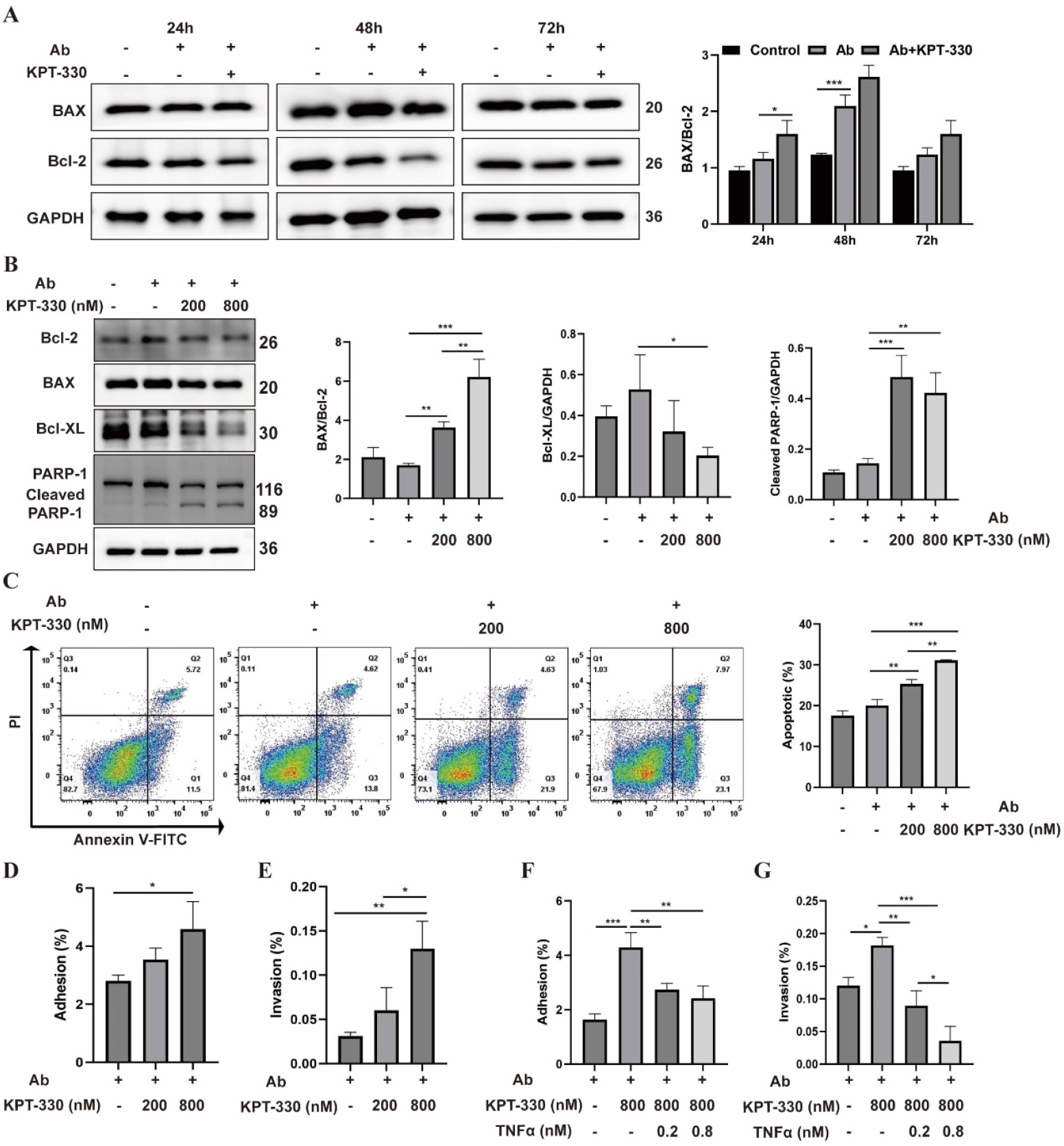
KPT-330 induces epithelial cell apoptosis by promoting bacterial adhesion and invasion. (A) Western blot analysis and grayscale statistics (n=3 each) assessing the expression of BAX and Bcl-2 proteins in lung tissues at 24 h, 48 h, and 72 h. (B) Western blot showing the expression of Bcl-2, BAX, Bcl-XL, and PARP-1 in Beas-2b cells. Beas-2b were treated with KPT-330 for 24 h prior to 6 h of Ab treatment. (C) Analysis of cell apoptosis by Annexin V-FITC/PI flow cytometry. KPT-330-pretreated (24 h) Beas-2b were harvested for flow cytometric analysis 6 h post-Ab treatment. (D, E) Effect of KPT-330 on the adhesion and invasion of Ab to Beas-2b cells with a Multiplicity of Infection (MOI) of 10:1. For the adhesion assay, Beas-2b cells were pre-treated with KPT-330 (200 nM or 800 nM) for 24 h then treated with Ab for another 2 h. For the invasion assay, cells were further treated with 100 μM gentamicin for 1 h to remove cell surface bacteria. (F, G) Effect of KPT-330 on the adhesion and invasion of Ab to Beas-2b cells in the presence of TNFα. KPT-330-pretreated Beas-2b cells were stimulated with TNFα (0.2 or 0.8 nM) for 6 h, then incubated with Ab (MOI 10:1) for another 2 h. Data are presented as mean ± SD. Differences between groups were compared using one-way ANOVA. *p < 0.05, **p < 0.01, ***p < 0.001.

As apoptotic cells are more susceptible to bacterial adhesion and invasion,^19-21^ we investigated whether KPT-330 influences these processes. Indeed, KPT-330 dose-dependently enhanced both Ab adhesion and invasion of Ab in Beas-2b cells (Fig. 3D and 3E). This enhanced bacterial adhesion and invasion were dose-dependently reversed by the addition of TNFα (Fig. 3F and 3G), likely through the activation of anti-apoptotic NFκB signaling.^22-24^ KPT-330 at physiological relevant doses had no direct effect on Ab proliferation or biofilm formation (Fig. S2), indicating that its effects were mediated through host changes. Collectively, these results demonstrate that during infection, KPT-330 induces apoptosis in lung epithelial cells and promotes bacterial adhesion/invasion, partly explaining for the observed higher bacterial load.

### Leukocyte recruitment is initially inhibited by KPT-330 and subsequently exaggerated

Leukocytes play a critical role in inflammation. We isolated total leukocytes from lung tissues, which are primarily recruited from the circulation during acute infection,^25, 26^ to determine the effects of Ab and KPT-330 (Fig. 4A). Results showed that Ab infection increased CD45+ leukocyte levels by 24 h, which remained largely unchanged by 72 h (Fig. 4B). KPT-330 suppressed this infiltration at 24 h (from 1.65% to 0.58%) but resulted in a significant rebound by 72 h (from 1.63% to 2.25%). In contrast to the declining trend of neutrophil in the Ab group, KPT-330-treated mice exhibited a continuous increment in neutrophil over time (Fig. 4C). Macrophage levels in the KPT-330 group followed a rise-and-fall pattern like the Ab group, but remained approximately two-fold higher at 72 h (Fig. 4D). These results suggest that KPT-330 suppresses inflammatory cell recruitment initially, but at later timepoints (48 h and 72 h), as the pharmacologic effect of KPT-330 diminished, a robust rebound infiltration occurred, likely triggered by the sustained bacterial load.

**Fig. 4.**
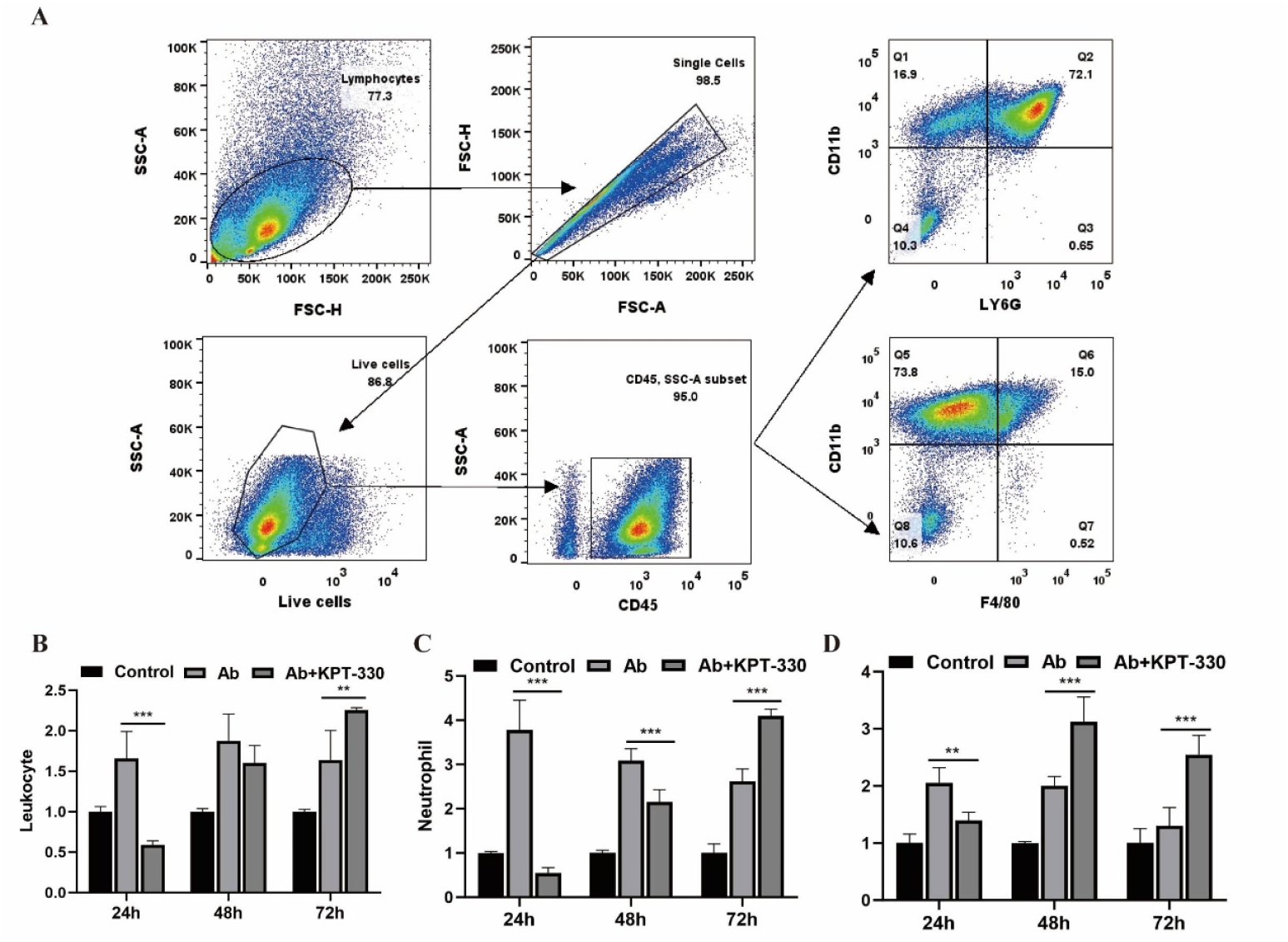
Leukocyte recruitment is initially inhibited by KPT-330 and subsequently exaggerated. (A) Sequential gating strategy employed to identify pulmonary immune cell subsets. Single-cell suspensions were prepared from lung tissues digested with collagenase/DNase I, followed by debris exclusion and dead cell removal. Immune cells were defined as the CD45^+^ population. Neutrophils were identified as the CD11b+Ly6G+ population, while macrophages were defined as CD11b+F4/80+ cells. (B) The proportion of leukocyte in mouse lung tissues (n=4 each). (C) The proportion of neutrophils in lung tissues (n=4 each). (D) The proportion of macrophages in lung tissues (n=4 each). All data were statistically compared to the control group. Data are presented as mean ± SD. Differences between groups were compared using unpaired Student’s t test. **p < 0.01, ***p < 0.001.

### NFκB signaling is inhibited at 24 h then activated at 48 h and 72 h in KPT-330-treated mice

NFκB signaling, a critical inflammatory pathway, was previously reported to be inhibited by KPT-330. To investigate its role in our model, we first used ELISA to measure the expression of key NFκB-controlled inflammatory cytokines IL-6, IL-1β, and TNFα, in the lung tissues and BALF. In the lung tissues, KPT-330-treated mice showed reduced IL-6/IL-1β levels at 24 h and reduced TNFα at 48 h, compared to Ab infection alone (Fig. 5A-C). By 72 h, however, this suppression was replaced by a significant increase in all three proteins. A slightly different kinetic pattern was observed in BALF, where KPT-330 treatment had minimal effect at 24 h but dramatically enhanced the levels of IL-6/IL-1β levels at 48 h and all three cytokines at 72 h (Fig. 5D-F). The protein level changes were transcriptionally regulated, as the qRT-PCR analysis showed a biphasic pattern of mRNA repression at 24 h followed by upregulation at 72 h (Fig. S3, Table S1).

**Fig. 5.**
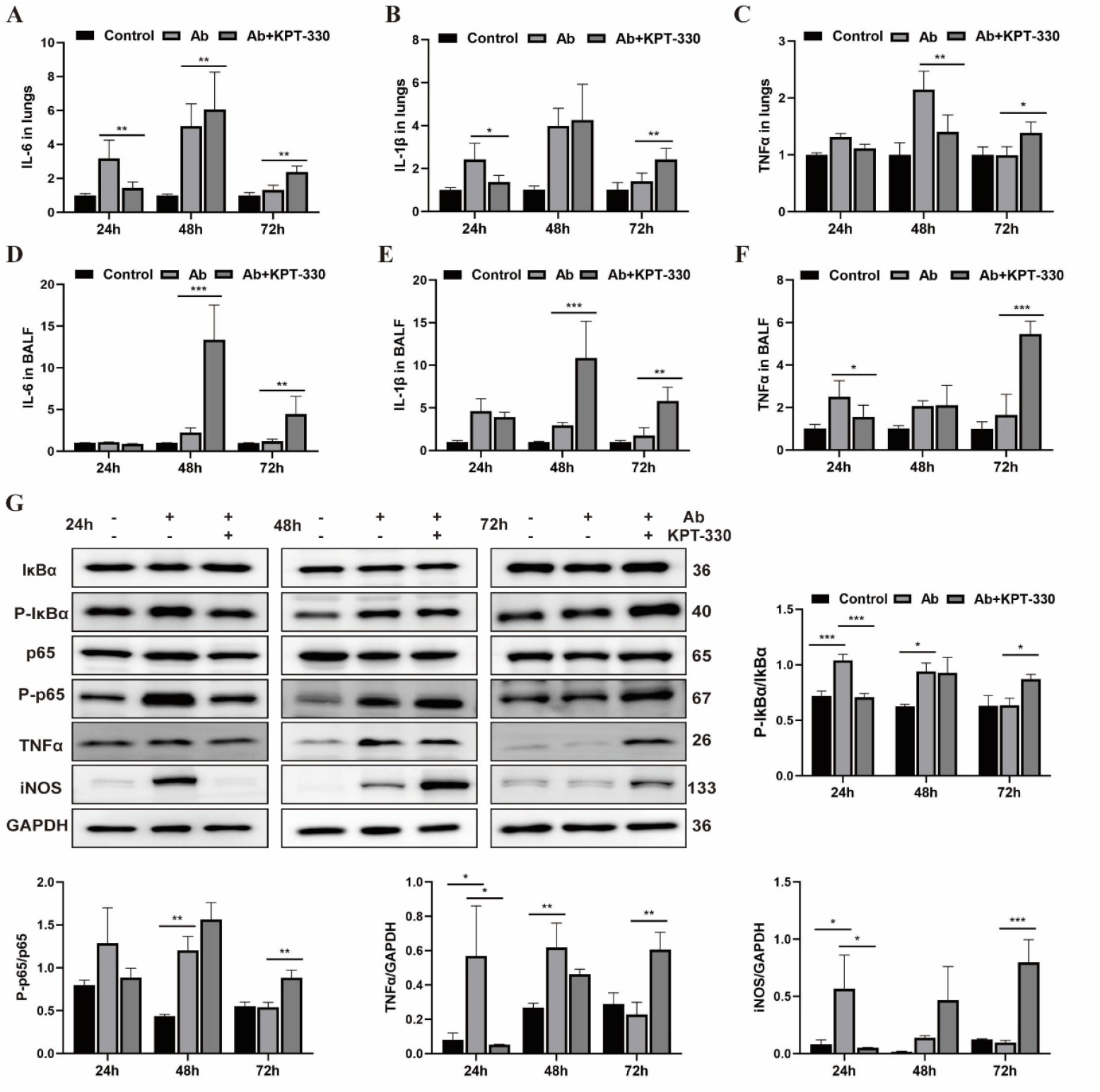
NFκB signaling is inhibited at 24 h then activated at 48 h and 72 h in KPT-330-treated mice. (A-C) Levels of IL-6, IL-1β, and TNFα in supernatants of homogenized mice lung tissues, quantified by ELISA at 24, 48, and 72 h post-stimulation. Data were normalized to the control group (n=5 each). (D-F) Levels of IL-6, IL-1β, and TNF-α in supernatants of BALF, quantified by ELISA at 24, 48, and 72 h post-stimulation. Data were normalized to the control group (n=5 each). (G) Western blot analysis of the expression of p65, p-p65, IκBα, P-IκBα, TNFα, and iNOS in the lung tissues of mice at 24 h, 48 h, and 72 h (n=3 each). Data are presented as mean ± SD. Differences between groups were compared using unpaired Student’s t test. *p < 0.05, **p < 0.01, ***p < 0.001.

We further characterized key NFκB pathway factors in lung tissues by Western blot. Phosphorylation of IκBα, the rate-limiting step for NFκB activation, was significantly reduced by KPT-330 at 24 h but enhanced at 72 h (Fig. 5G). A parallel biphasic pattern (inhibition and activation) was observed for the phosphorylation of p65, which is critical for NFκB transcriptional activity. Consistent with these signaling dynamics, expression of the downstream transcribed effectors TNFα and iNOS was initially suppressed and subsequently upregulated in KPT-330-treated mice (Fig. 5G). Collectively, these data demonstrate that KPT-330-treated mice display a biphasic regulation of the NF-κB pathway, which provides a mechanistic explanation for the biphasic leukocyte recruitment observed earlier.

### KPT-330 blocks the nuclear export of p65 and IκBα in Beas-2b cells

Both p65 and IκBα contain functional nuclear export signals (NESs) that are crucial for their nuclear export.^27, 28^ Here, we found that KPT-330 treatment blocked the nuclear export of both proteins in Beas-2b cells (Fig. 6A-D), consistent with its known function. Although nuclear p65 activates transcription, nuclear IκBα potently inhibits NFκB signaling by stripping it from DNA.^29^ Western blot analysis revealed that KPT-330 reduced IκBα phosphorylation (enhancing its binding/inhibition to NFκB) and also suppressed p65 phosphorylation (rendering it less transcriptionally active), leading to decreased TNFα transcription in Beas-2b cells (Fig. 6E). The XPO1 levels is also reduced by KPT-330, consistent with previous reports.^30, 31^ These results provide a mechanistic basis for the transient anti-inflammatory and lung-protective effect of KPT-330.

**Fig. 6.**
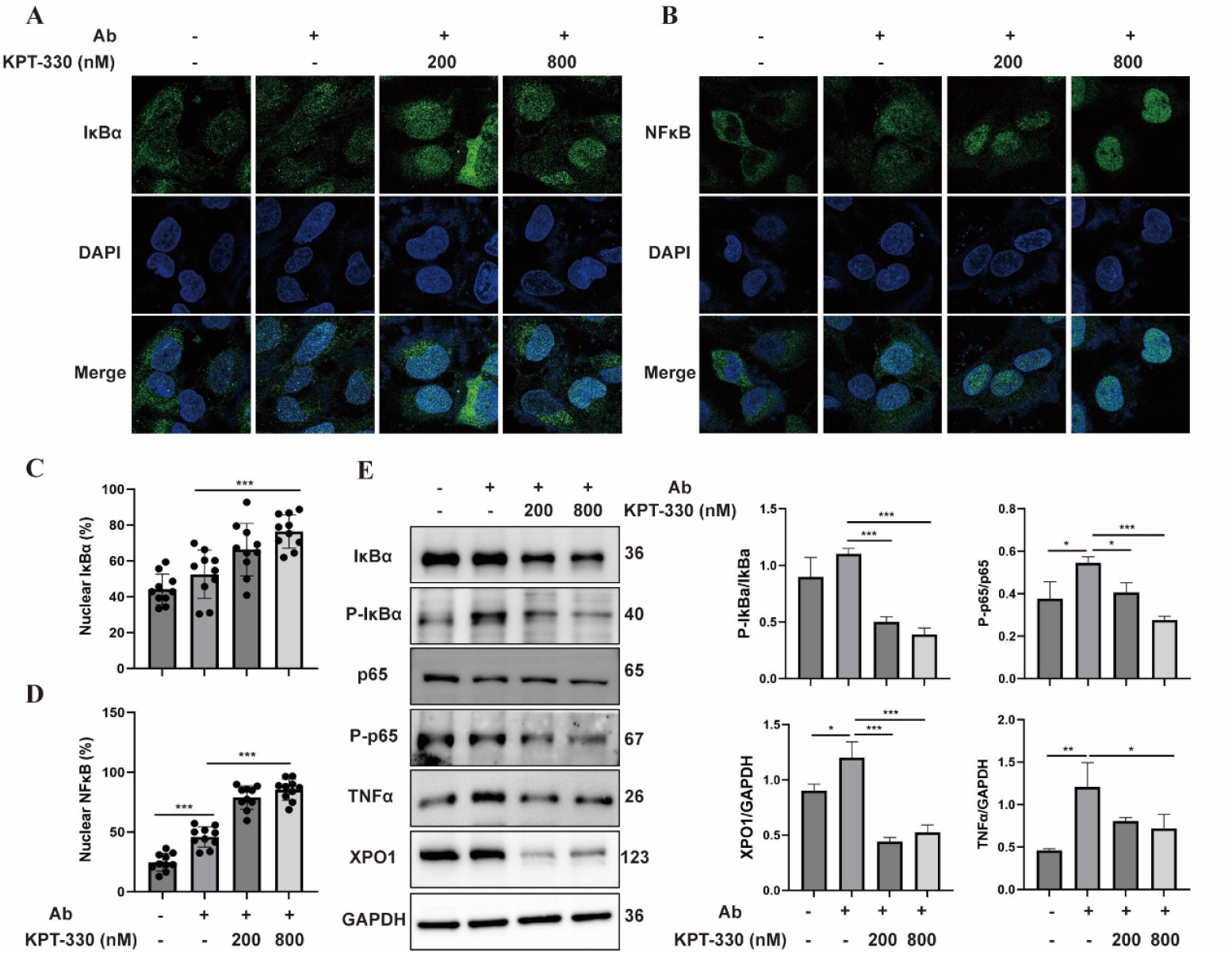
KPT-330 blocks the nuclear export of p65 and IκBα in Beas-2b cells. (A, B) Immunofluorescence images of IκBα and p65 in Beas-2b cells pre-treated with different concentrations of KPT-330 for 24 h and Ab (MOI 10:1) for another 6 h. (C, D) Quantification and statistical analysis of the nuclear IκBα and p65 percentages (nuclear/total). (E) Western blot analysis of the expression of IκBα, P-IκBα, p65, P-p65, TNFα, and XPO1 in KPT-330- and Ab-treated Beas-2b cells. Data are presented as mean ± SD. Differences between groups were compared using one-way ANOVA. *p < 0.05, **p < 0.01, ***p < 0.001.

### Polymyxin E but not dexamethasone improves the survival of KPT-330-treated mice

We further investigated whether KPT-330-induced lethality could be reversed with adjunct therapies that combat the rise in bacteria load and inflammation. Results showed that the antibiotic drug polymyxin E (5 mg/kg) significantly enhanced the survival of KPT-330-treated mice (Fig. 7A). Polymyxin E treatment slightly reduced lung wet-to-dry (W/D) ratio and lung index at 24 h, although these differences were not statistically significant (Fig. 7B and 7C). Furthermore, polymyxin E dramatically reduced the bacterial load in the lung at 24h by 9-fold (Fig. 7D). Albeit not statistically significant, bacterial levels in blood, liver, spleen, and kidney were also reduced (Fig. S4). Using Beas-2b cells, we demonstrated that Ab potently induced cell membrane leakage as measured by LDH release, which was mildly potentiated by KPT-330 (Fig. 7E), revealing a mechanism for Ab-induced lung damage. These results confirm that adjunct antibiotics such as polymyxin E mitigates lethality from KPT-330 monotherapy by reducing bacterial load and associated damage (Fig. 7F).

**Fig. 7.**
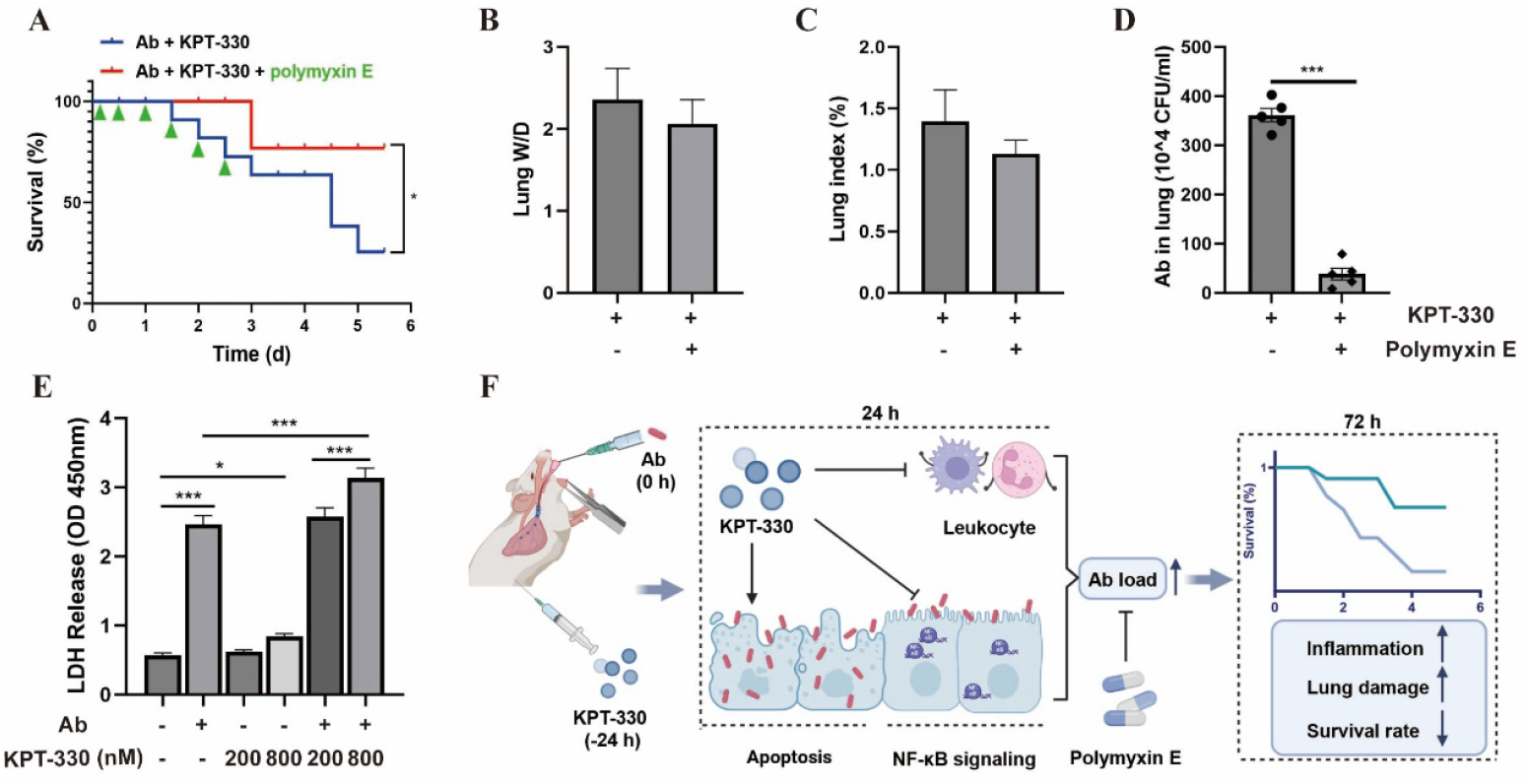
Polymyxin E improves the survival of mice treated with KPT-330. (A) Survival rates of KPT-330-treated Ab-infected mice treated with or without polymyxin E (5 mg/kg). Polymyxin E (every 12 h for 6 times) was intraperitoneally administered 2 h following intratracheal Ab injection (n=9 each). (B) Wet-to-dry ratio of the lungs before and after administration of polymyxin E (n=5 each). (C) Lung index in mice before and after administration of polymyxin E (n=5 each). (D) Bacterial load in lungs before and after administration of polymyxin E (n=5 each). (E) LDH release in Beas-2b cells after 10 h treatment with KPT-330 (200 nM or 800 nM) and/or Ab (MOI 1:10). (F) A schematic diagram showing the Janus face of KPT-330 in pneumonia caused by Ab infection. At 24 h post-infection, KPT-330 suppressed NF-κB signaling and leukocyte recruitment while promoting lung epithelial apoptosis, consequently leading to a high bacterial load. By 72 h, mice suffered reduced survival due to the consequent severe inflammation and lung damage, a lethality that was preventable by polymyxin E treatment. Data are presented as mean ± SD. *p < 0.05, **p < 0.01, ***p < 0.001.

In contrast, combining KPT-330 with the anti-inflammatory drug dexamethasone did not significantly improve survival (Fig. S5), although it inhibited lung W/D ratio and lung index at 48 h (Fig. S6). Dexamethasone significantly increased the bacterial load in the lung, blood, liver, and kidney (Fig. S7), consistent with its immune-suppressive characteristics. Although dexamethasone increased the bacterial load, it did not further reduce survival compared to the KPT-treated group, suggesting that its anti-inflammatory effect provided certain survival benefits that counterbalanced the detriment of enhanced bacterial growth.

## Discussion

In this work, we dissect the time-dependent effect of KPT-330 in a murine model of Ab pneumonia. Our findings reveal a critical paradox: although KPT-330 initially attenuated lung inflammation and damage, it ultimately predisposed the host to severe, lethal infection by compromising bacterial clearance and increasing epithelial vulnerability.

A key finding is that KPT-330 markedly increased the bacterial burden in mice. This effect was attributed to two primary reasons: 1) KPT-330 suppressed immune cell recruitment as well as anti-bacterial inflammatory signaling, permitting uncontrolled bacterial replication in the lungs, and 2) KPT-330 induced epithelial apoptosis, thereby promoting bacterial adhesion and invasions, and potentially facilitating system dissemination. The increasing bacterial load, together with the ability of Ab to damage lung epithelial cells, would impair pulmonary function, leading to increased mortality of KPT-330-treated mice. The finding that mortality was effectively reversed by adjunct therapy with polymyxin E confirms that failure of timely bacterial clearance was the primary driver of the lethal outcome.

Consistent with previous studies,^32, 33^ we demonstrated that KPT-330 also suppressed the nuclear export of IκBα in Beas-2b cells, resulting in potent suppression of NFκB. This mechanism underpinned the transient anti-inflammatory and lung-protective effect observed at 24 h post-infection. Furthermore, our results demonstrate a strong suppressive effect of KPT-330 on neutrophils, a finding consistent with the drug’s known clinical side effect of neutropenia.^6, 34, 35^ This immunosuppression permitted a persistently high bacterial burden, which subsequently triggered a robust immune rebound after 48 h and 72 h. This “immune rebound” is not merely a quantitative increase in leukocytes, but represents a dysregulated and potentially harmful inflammatory response. This explains the observation that dexamethasone, while further increasing the bacterial load due to its immunosuppressive effects, did not worsen survival compared to KPT-330 alone. Therefore, prolonged and heightened inflammatory response likely also contributed to the poor survival in the KPT-330-treated group.

Our study carries important clinical implications, as KPT-330 can exacerbate the susceptibility of cancer patients to opportunistic bacterial infections. Given that most, if not all, patients eligible for KPT-330 have undergone multiple lines of prior therapy and exhibit substantially compromised immune function,^14, 36^ our work support the integration of prophylactic antibiotics into treatment for patients receiving KPT-330. Similar prophylactic strategies have been successfully implemented on other immunosuppressive targeted therapies such as Bruton’s tyrosine kinase (BTK) inhibitors.^37, 38^ While polymyxin E was effective in our model, real-world prophylaxis may consider broad-spectrum agents to cover a wider range of potential pathogens. The immune-suppressive effects of KPT-330, while detrimental in the context of infection, may contribute to its efficacy in hematologic malignancies by modulating the tumor microenvironment.^39^ This duality underscores the critical need for personalized risk-benefit assessments and vigilant monitoring for infections during XPO1 inhibitor therapy.

While our study identifies the risk of KPT-330 in Gram-negative pneumonia, the drug-induced immune suppression suggests that this susceptibility may extend beyond this class of pathogens. Consequently, the potential for a broader infection risk, including from fungi and viruses, warrants increased clinical vigilance and further investigation. Furthermore, as KPT-330 is clinically administered once or twice per week, our pre-infection model is physiologically relevant. However, further studies are needed to determine whether post-infection administration yields a similar outcome. Finally, the observed reduction in bacterial adhesion and invasion by TNFα merits future investigation as a potential adjunctive therapy.

## Acknowledgments

This work was supported by the National Natural Science Foundation of China [Grant Number 82273850 to Q.S.]. We also thank Dr. Yuling Li (University of Electronic Science and Technology of China) for helpful scientific discussions.

## Author contributions

Conceptualization: QY, QS

Methodology: QY, QS

Investigation: QY

Visualization: QY

Funding acquisition: LG, QS

Project administration: LG, QS

Supervision: LG, QS

Writing - QY, QS

Writing - review & editing: QY, LG, QS

## Data and materials availability

All data are available in the main text or the supplementary materials.

## Materials and Methods

### Animals and infection model

*Acinetobacter baumannii* (strain ATCC 17978) was cultured in LB medium (Solarbio) at 37°C with shaking at 250 rpm. C57BL/6 male mice (6 weeks old, 18 -20 g) were purchased from SPF Biotechnology Co., Ltd. and housed under a 12 h light/dark cycle at 23 ± 1°C with free access to food and water. Mice were randomly assigned to three groups: (1) vehicle control, (2) Ab infection alone (Ab), or (3) Ab infection plus KPT-330 (Ab + KPT-330). KPT-330 (selinexor; Kaiwei Chemical) was administered via intraperitoneal injection (20 mg/kg) 24 h before bacterial inoculation. For infection, mice were anesthetized with avertin (Mreda) and subjected to intratracheal instillation of Ab (8 × 10^7 CFU) to induce pneumonia. Lung tissues and bronchoalveolar lavage fluid (BALF) were collected at 24 h, 48 h, and 72 h post-infection. To quantify the wet-to-dry weight ratio, freshly harvested lungs were immediately weighed (wet weight), incubated at 65°C for 24 h, and then weighed again (dry weight). The same regions of lung tissues were fixed in 10% buffered formalin, embedded in paraffin, sectioned, and stained with hematoxylin and eosin (H&E). Morphological changes were assessed by light microscopy. The remaining lung tissue was homogenized for bacterial load quantification or snap-frozen and stored at -80°C. All animal experiments were approved by the Animal Ethics Committee of Sichuan Provincial People’s Hospital (approval no. 2025-418) and conducted in compliance with relevant ethical regulations.

### Survival analysis

To assess the effect of KPT-330 on survival, mice were treated intraperitoneally with KPT-330 or vehicle 24 h before intratracheal infection with Ab. In separate therapeutic intervention experiments, infected mice received intraperitoneal polymyxin E (5 mg/kg) beginning 2 h post-infection, or dexamethasone beginning 24 h post-infection. Both polymyxin E and dexamethasone were administered every 12 h for 3 days. Mouse survival was monitored every 12 h for 5 days, after which all surviving animals were euthanized.

### Bacteriological culture

After avertin anesthesia, mice were euthanized for sample collection. Orbital blood was collected and plated directly onto LB agar. BALF was obtained by instilling and aspirating 0.5 mL of PBS into the lungs three times. Lung, spleen, liver, and kidney tissues were homogenized in PBS. BALF and tissue homogenates were serially diluted, plated on LB agar, and incubated overnight at 37°C.

### Cytokine measurement

Concentrations of TNFα, IL-1β, and IL-6 in BALF and lung homogenates were measured using commercial ELISA kits from Ruixin Industrial Co., Ltd. (Quanzhou, China), according to the manufacturer’s instructions.

### Western blotting

Lung tissues were homogenized in lysis buffer containing protease and phosphatase inhibitors and lysed on ice for 30 min. Lysates were centrifuged at 12,000 × g for 10 min at 4°C. Protein concentrations in the supernatants were quantified, and equal amounts of protein were denatured in Laemmli buffer. Proteins were separated by SDS-PAGE and transferred to PVDF membranes (Merck KGaA). After blocking with 5% BSA, membranes were probed overnight at 4°C with the following primary antibodies: anti-p65 (Servicebio, GB11997-50), anti -P-p65 (Proteintech, 82335-1-RR), anti-IκBα (HuaBio, ET1603-6), anti-p-IκBα (HuaBio, HA722770), anti-TNF-α (Affinity, AF7014), anti-iNOS (Selleckchem, F0177), anti-XPO1 (Absin, abs115104), anti-Bcl-2 (Zenbio, 381702), anti-Bcl-XL (Zenbio, R23603), anti-PARP1 (Proteintech, 66520-1-Ig), anti-BAX (Selleckchem, F0037), and anti-GAPDH (Proteintech, 60004-1-Ig). Membranes were then incubated with HRP-conjugated secondary antibodies, and proteins were detected using an ECL kit (Yeasen Biotechnology).

### Flow cytometry

Single-cell suspensions were prepared from perfused murine lungs. Tissues were minced and digested at 37°C with collagenase I (5 mg/ml) and DNase I (1 mg/ml) (Solarbio) for 40 min. The resulting suspension was passed through a 70-μm strainer, and erythrocytes were lysed.

Cells were washed, resuspended in flow cytometry buffer (PBS with 5% FBS), and counted. For staining, cells were incubated with a fixable viability dye (eFluor 450; Invitrogen, 65-0863-14) and the following antibodies: anti-CD45 (ABclonal, A23709), anti-CD11b (ABclonal, 6950001153), anti-Ly6G (PE-Cy™7; Elabscience, I-AB-F1108H), and anti-F4/80 (ABclonal, 6950000458). Beas-2b cells were harvested using EDTA-free trypsin, washed, and stained with Annexin V-FITC and propidium iodide (PI) (Beyotime Biotechnology) according to the manufacturer’s protocol. Data were acquired on a BD FACSymphony A1 flow cytometer and analyzed using FlowJo software.

### Bacterial adhesion and invasion assays

The adhesion and invasion experiments were adapted from a previously described protocol.^40^ Human bronchial epithelial Beas-2b cells (Servicebio) were cultured in DMEM supplemented with 10% FBS and 1% penicillin-streptomycin at 37°C with 5% CO_2_. Cells were pretreated with KPT-330 for 24 h and then challenged with Ab at a MOI of 10:1 for 2 h. For adhesion, infected cells were washed with PBS and lysed with 0.5% Triton X-100. For invasion, extracellular bacteria were killed by a 1 h gentamicin (100 µM) treatment post-infection before lysis. Lysates were serially diluted and plated for bacterial enumeration.

### Immunofluorescence

Beas-2b cells grown on coverslips were fixed with 4% PFA, permeabilized with 0.1% Triton X-100, and blocked with 5% BSA. Cells were incubated with primary antibodies overnight at 4°C, followed by fluorescent secondary antibodies for 1 h at room temperature. Nuclei were stained with DAPI. Images were captured using a Zeiss fluorescence microscope.

### Statistical analysis

Data are presented as mean ± SD from at least three independent samples. Statistical analyses were performed using GraphPad Prism software. An unpaired, two-tailed Student’s t test was used for comparisons between two groups. One-way ANOVA was used for comparisons among multiple groups. A p value of less than 0.05 was considered statistically significant.

